# Spatiotemporal Dynamics of Orthographic and Lexical Processing in the Ventral Visual Pathway

**DOI:** 10.1101/2020.02.18.955039

**Authors:** Oscar Woolnough, Cristian Donos, Patrick S. Rollo, Kiefer J. Forseth, Yair Lakretz, Nathan E. Crone, Simon Fischer-Baum, Stanislas Dehaene, Nitin Tandon

**Affiliations:** Vivian L. Smith Department of Neurosurgery, McGovern Medical School at UT Health Houston, TX, 77030, USA; Texas Institute for Restorative Neurotechnologies, University of Texas Health Science Center at Houston, Houston, TX, 77030, USA; Faculty of Physics, University of Bucharest, 050663, Bucharest, Romania; Cognitive Neuroimaging Unit CEA, INSERM, NeuroSpin Center, Université Paris-Sud and Université Paris-Saclay, 91191, Gif-sur-Yvette, France; Department of Neurology, Johns Hopkins Medical Center, Baltimore, MD, 21287, USA; Department of Psychological Sciences, Rice University, Houston, TX, 77005, USA; Collège de France, 11 Place Marcelin Berthelot, 75005, Paris, France; Memorial Hermann Hospital, Texas Medical Center, Houston, TX, 77030, USA

**Author notes:** Corresponding Author: Nitin Tandon.

## Abstract

Reading is a rapid, distributed process that engages multiple components of the ventral visual stream. However, the neural constituents and their interactions that allow us to identify written words are not well understood. Using direct intracranial recordings in a large cohort of humans, we comprehensively isolated the spatiotemporal dynamics of visual word recognition across the entire left ventral occipitotemporal cortex. The mid-fusiform cortex is the first region that is sensitive to word identity and to both sub-lexical and lexical frequencies. Its activation, response latency and amplitude, are highly dependent on the statistics of natural language. Information about lexicality and word frequency propagates posteriorly from this region to traditional visual word form regions and to earlier visual cortex. This unique sensitivity of mid-fusiform cortex to the lexical characteristics of written words points to its central role as an orthographic lexicon, which accesses the long-term memory representations of visual word forms.

## Introduction

Reading is foundational to modern civilization, yet it is a very recently acquired cultural skill. The substrates that allow us to fluently convert orthography to semantic information are not well understood. The ventral occipitotemporal cortex (vOTC) has been thought to decipher orthographic information of various levels of complexity, in a posterior to anterior gradient (Binder et al., 2006; Dehaene et al., 2005; Vinckier et al., 2007). Its greater activation by pseudowords (Kronbichler et al., 2004), and low frequency words (Graves et al., 2010; Kronbichler et al., 2004; Schuster et al., 2016; White et al., 2019), points to its crucial role in word identity. Orthographic representations are thought to be organized hierarchically in the vOTC with bottom-up processes culminating in the visual word form area (VWFA) (Dehaene and Cohen, 2011; Dehaene et al., 2002, 2005), a word selective region that is also involved in sub-lexical processing. Recent evidence suggests however that this view may be oversimplified and the VWFA may instead be comprised of multiple functionally distinct patches engaged in perceptual (orthographic vs. pattern), sub-lexical (word vs. non-word) or lexical (whole word level) derivations (Lerma-Usabiaga et al., 2018; White et al., 2019). More importantly, it remains unclear what roles specific components of the ventral stream play in the integration of bottom-up information (Dehaene and Cohen, 2011; Dehaene et al., 2002, 2005) with top-down influences from higher language regions (Kay and Yeatman, 2017; Pammer et al., 2004; Price and Devlin, 2003, 2011; Song et al., 2010; Starrfelt and Gerlach, 2007; Whaley et al., 2016; White et al., 2019; Woodhead et al., 2014) to enable rapid orthographic-lexical-semantic transformations.

While most of our knowledge of the cortical architecture of reading arises from functional MRI, the rapid speed of reading demands that we use methods with very high spatiotemporal resolution to study these processes. To this end, we used recordings in 35 individuals with 784 intracranial electrodes, to comprehensively characterize the spatial organization and functional roles of orthographic and lexical regions across the ventral visual pathway during sub-lexical and lexical processes. Given their construction, these two tasks, performed in the same cohort, tap into varying levels of attentional modulation of orthographic processing. Specifically, we isolated functionally distinct regions across the vOTC that are highly sensitive to the structure and statistics of natural language at multiple stages of orthographic processing.

## Results

Patients participated in sub-lexical and lexical tasks specifically designed to disambiguate the roles of sub-regions and top-down attentional modulation within the vOTC. In the sub-lexical task, patients viewed strings of false font characters (FF), infrequent letters (IL), frequent letters (FL), frequent bigrams (BG), frequent quadrigrams (QG) or words (W) while detecting a non-letter target (Figure 1a,b). In the lexical task, patients attended to regular sentences, word lists or jabberwocky sentences, all presented in rapid serial visual presentation format, followed by a forced choice decision of presented vs non presented words (Figure 1c,d).

**Figure 1:**
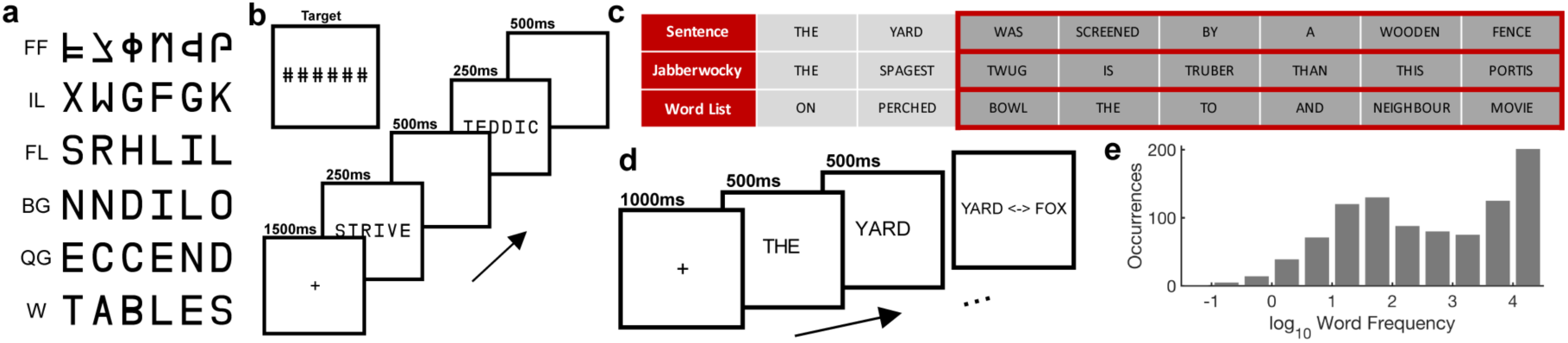
Experimental design of the sub-lexical and lexical tasks. (a) Example stimuli from each of the six sub-lexical stimulus categories. FF: False Font, IL: Infrequent Letters, FL: Frequent Letters, BG: Frequent Bigrams, QG: Frequent Quadrigrams, W: Words. (b) Schematic representation of the sub-lexical stimulus presentation. (c) Example stimuli from the three experimental conditions of the lexical task highlighting the words used for subsequent analyses (Words 3 to 8). (d) Schematic representation of the lexical stimulus presentation. (e) Histogram of log_10_ word frequency for the sentence stimuli. A frequency of 1 represents 10 instances per million words and 4 meaning 10,000 instances per million words

Word responsive electrodes (defined as >20% gamma band activation above baseline) were seen across the entire vOTC from the occipital pole to mid-fusiform cortex in the left, language-dominant hemisphere, and only in the occipital pole of the right hemisphere (Figure 2a; Figure S1). Thus, we constrain all our analysis to the left hemisphere.

**Figure 2:**
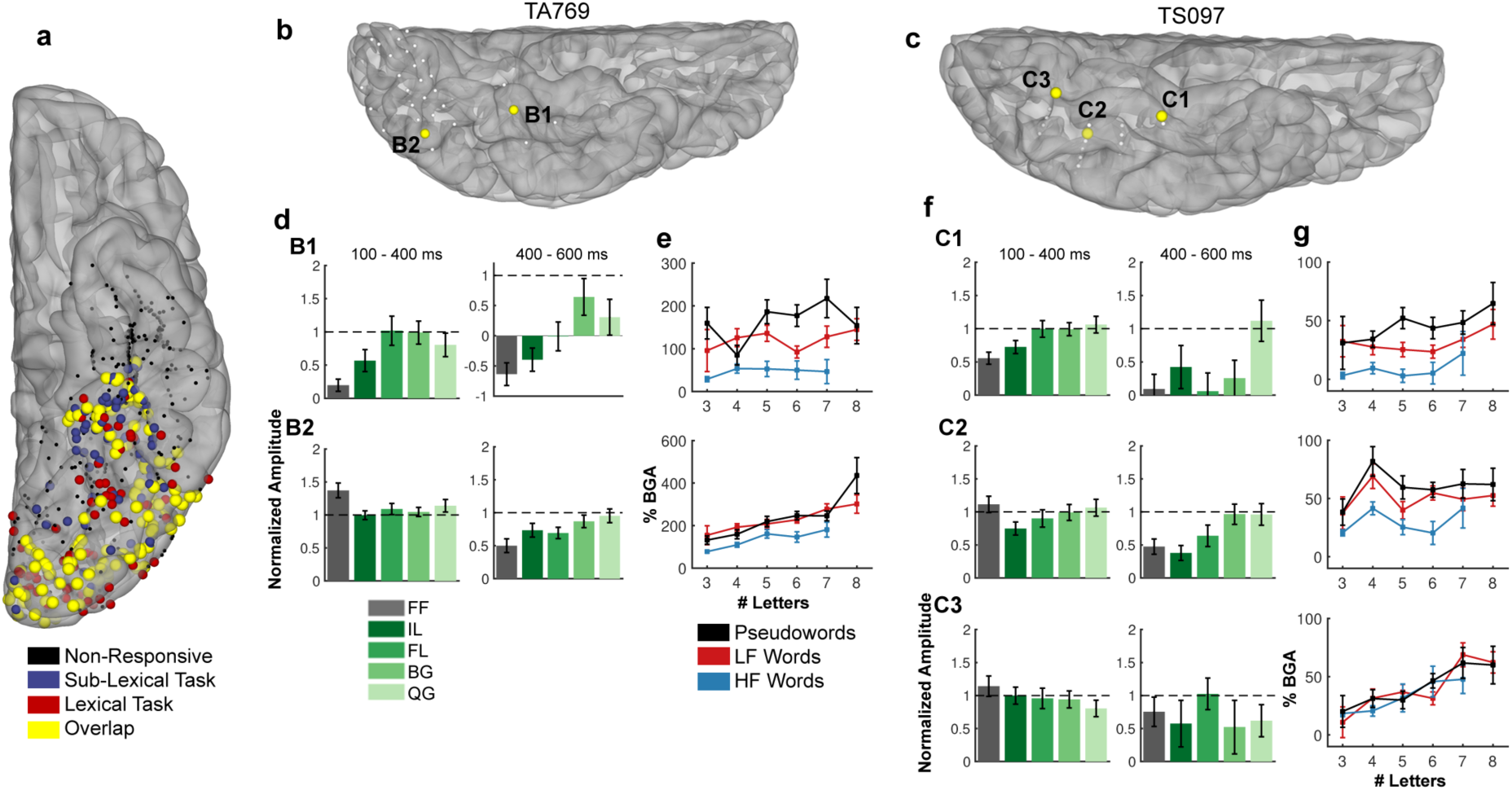
Population activation map and single patient activations. (a) Locations of all electrodes within the left vOTC ROI that were responsive to real words (>20% activation over baseline) in patients that performed the sub-lexical task (blue), the lexical task (red), those that performed both tasks (yellow) and those electrodes that were not responsive (black). (b,c) Location of electrodes within single subjects, demonstrating posterior-to-anterior changes in responses to each task. (d,f) Word normalized selectivity profiles in the sub-lexical task at early (100-400ms; left) and late (400-600ms; right) time points; (e,g) plots of broadband gamma activity (BGA; 70 −150 Hz) sensitivity to length, frequency (high frequency (HF) and low frequency (LF) words), and lexical status.

### Orthographic Processing in vOTC

We characterized activity in early (100-400 ms; presumably reflecting automatic visual processes (Kadipasaoglu et al., 2016)) and late (400-600ms) windows. Based on our earlier work (Forseth et al., 2018; Kadipasaoglu et al., 2016) and on other studies detailing the roles of these regions (Lerma-Usabiaga et al., 2018; White et al., 2019), we also separated the vOTC into anterior and posterior sites (y = −40 mm). Anterior sites in the vOTC were less responsive to FF and IL stimuli than to words throughout, more so in the later time windows when this region stayed responsive to words for longer than all other stimuli (Figure 2d,f). Further, these anterior regions showed a marked distinction between high and low frequency words (derived from an American English language corpus, SUBTLEXus (Brysbaert and New, 2009)) and a smaller effect of word length (Figure 2e,g).

Posterior regions responded the most to false fonts and did not distinguish between other non-word stimuli in early time windows (100-400 ms), however at later time points (400-600 ms) sensitivity to sub-lexical complexity was seen. In the lexical task, these posterior sites were sensitive to word length and less so to word frequency.

To evaluate these effects at the population level, we performed a mixed-effects, multilevel analysis (MEMA) of broadband gamma (70-150 Hz) activation between 100-400 ms, in grouped normalized 3D stereotactic space. This analysis is specifically designed to account for sampling variations and to minimize effects of outliers (Argall et al., 2006; Conner et al., 2014; Esposito et al., 2013; Fischl et al., 1999; Forseth et al., 2018; Miller et al., 2007; Saad and Reynolds, 2012; Woolnough et al., 2019). This MEMA map showed that written words activated the left vOTC from occipital pole to mid-fusiform cortex (Figure 3a). We then used this map to delineate regions showing preferential activation for words compared to non-word stimuli (Figure 3b). A clear posterior-to-anterior transition - from occipital cortex to mid-fusiform gyrus - was observed. We again noted that in mid-fusiform cortex, responses to words were predominant, it distinguished between IL stimuli and real words but it did not show substantial difference between words and word-like stimuli (FL, BG and QG). This selectivity pattern was reversed in posterior occipitotemporal cortex which was more active for FF stimuli than for words.

**Figure 3:**
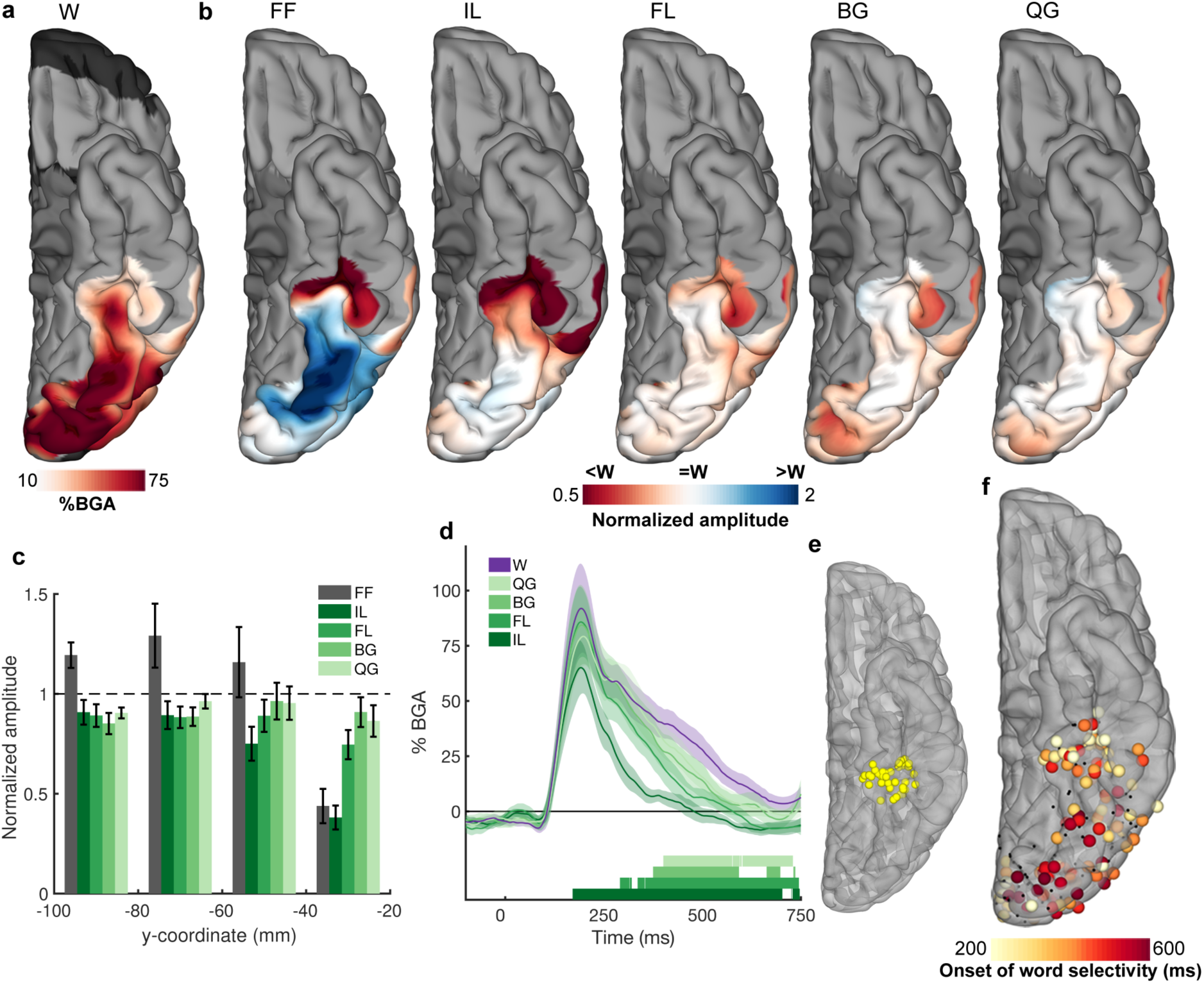
Spatial mapping of selectivity to hierarchical orthographic stimuli in the sub-lexical task. (a) MEMA activation map showing the regions of significant activation to the real word stimuli in the sub-lexical task (100 – 400 ms). This activation map was used as a mask and a normalization factor for the activations of the non-word stimuli (b). Normalized amplitude maps showing regions with preferential activation to words (red) or non-words (blue). (c) Electrode selectivity profiles grouped every 20 mm along the antero-posterior axis in Talairach space. (d) Contrasts of the lettered non-words against words, within mid-fusiform cortex (e), showing latency differences between when each non-word category can be distinguished from words. Colored bars under the plots represent regions of significant difference from words (q<0.05). (f) Spatial map of the initial timing of significant word selectivity. Electrodes that did not reach significance shown in black.

To further characterize this spatial gradient, we plotted responses as a function of electrode location along the y axis in Talairach space, there was a larger response to FF from −100 to −60 mm, while other non-lexical stimuli (IL, FL, BG and QG) led to a similar response to words (Figure 3c). Between −60 and −40 mm, the response to IL and, to a lesser extent, FL stimuli was significantly less than the response to words. Around −40 mm in the antero-posterior axis in Talairach space, a distinct transition occurred: the response to FF and IL collapsed to a low level - the response became fully selective for words and word-like stimuli (BG and QG).

A 4D representation of the evolution of the functional selectivity of the vOTC, (Video 1, Video 2) clearly illustrates the primacy of the mid-fusiform cortex in word identification. When looking within mid-fusiform cortex (39 electrodes, 13 patients) we also see latency differences between when each of the non-word classes can be distinguished from words (Figure 3d, Video 2). With increasing sub-lexical structure, the latency of word/non-word distinction increases.

This 4D representation also reveals that at later time periods (>300 ms) an anterior-to-posterior propagation of word selectivity occurs - posterior regions show greater sustained activity for words rather than non-words. To further elaborate this antero-posterior spread of word selectivity, we calculated the sensitivity index (d-prime for words vs all non-word stimuli) over time at each electrode in vOTC to find the earliest point where responses for real vs non-words separated (Figure 3e; p < 0.01 for >50ms). Again, the mid-fusiform cortex showed the earliest word selective response (∼250 ms) and this selectivity then progressed posteriorly to occipital pole (∼500 ms). A correlation of the latency of d-prime significance with the y axis further quantifies this anterior-to-posterior gradient (r = −0.33, p < 0.001).

In summary, word selective responses in vOTC during passive viewing are seen earliest in mid-fusiform cortex. This selectivity then spreads posteriorly to earlier visual regions such as posterolateral vOTC, which while active early, demonstrate word selectivity late.

### Lexical Processing in Mid-Fusiform Cortex

Next, we sought to examine how this spatiotemporal lexical response pattern relates to higher order processes, such as sentence reading, that engage the entire reading network. Lexical contrasts (MEMA) between high vs. low frequency words and real vs. pseudowords, of gamma activity between 100–400 ms after each word, revealed two significant clusters consistent across both contrasts – the mid-fusiform cortex and lateral occipitotemporal gyrus (Figure 4a,b). In this task, where words were attended, we saw an *inversion* of the word vs non-word selectivity seen in the previous, passive-viewing sub-lexical task - pseudowords showed greater activation than real words (Dehaene and Cohen, 2011; Kronbichler et al., 2004), pointing to the role of top-down attentional modulation of activity in this region

**Figure 4:**
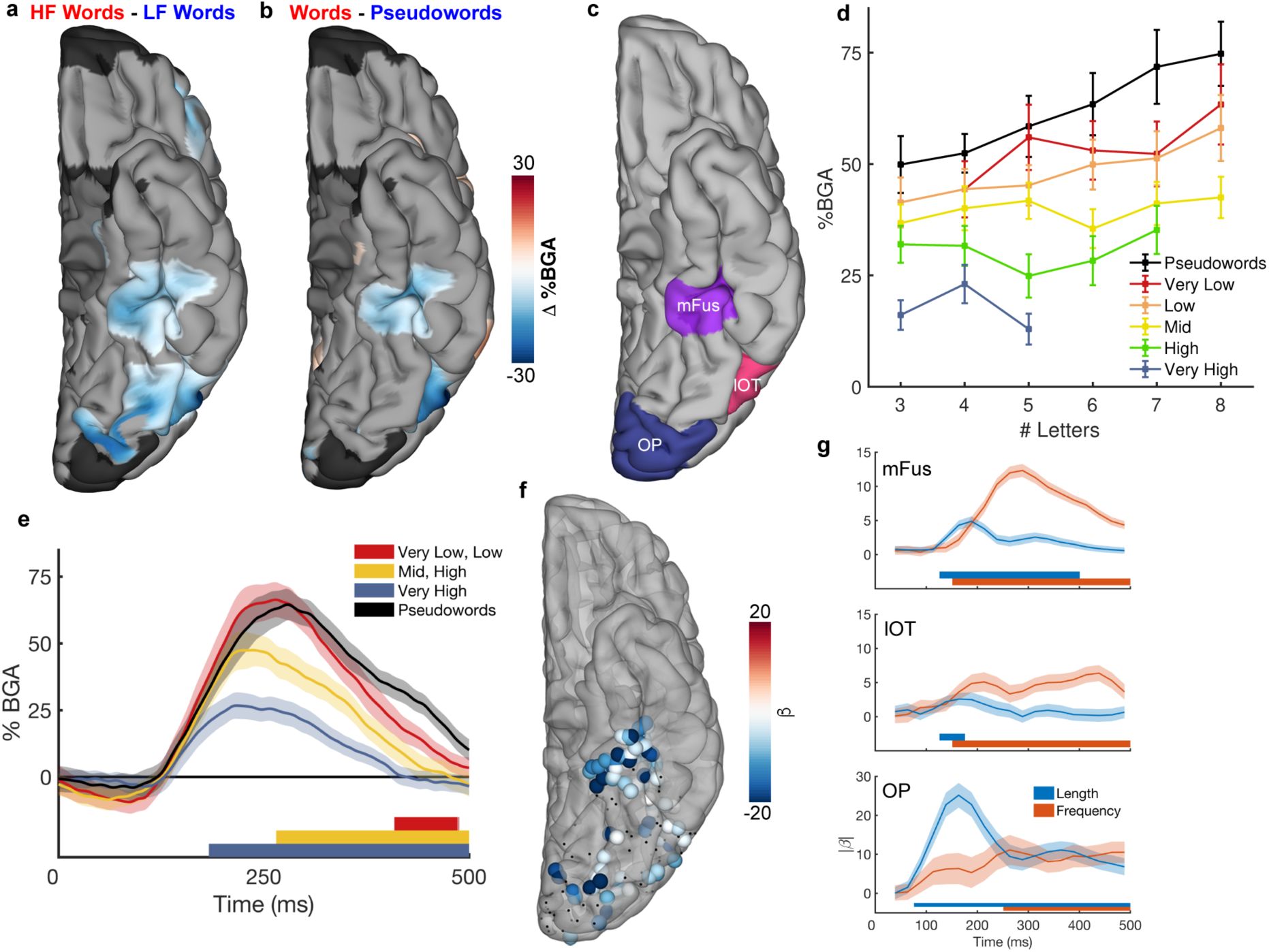
Spatiotemporal mapping of frequency and lexicality effects in the lexical task. Contrast MEMA results for (a) high frequency (HF; f > 2.5) vs low frequency (LF; f < 1.5) words from the sentence condition and (b) real words vs pseudowords (content words from sentences vs content words from Jabberwocky). Words were matched for length between conditions within each contrast. (c) ROI definitions. mFus: Mid-fusiform cortex, lOT: Lateral occipitotemporal cortex, OP: Occipital Pole. (d) Mid-fusiform cortex activation to real words from the sentence condition separated by word frequency and length. (e) Contrasts of different frequency words against pseudowords, within mid-fusiform cortex, showing latency differences between when each word frequency band can be distinguished from pseudowords. Colored bars under the plots represent regions of significant difference from pseudowords (q<0.05). (f) Individual electrodes showing significant modulation of gamma activity by word frequency. Electrodes that did not reach significance shown in black. (g) Time courses of length and frequency sensitivity within the three tested ROIs.

To quantify the relative sensitivity of mid-fusiform cortex (49 electrodes, 15 patients; Figure 4c) to word frequency and word length, we utilized a linear mixed effects (LME) model with fixed effects modelling word length and log word frequency (Figure 4d). A large proportion of the variance of this region’s activity (r^2^ = 0.73), is explained by word frequency (β = −8.5, p < 10^−40^), word length has a much smaller effect (β = 1.8, p < 10^− 4^). The interaction between these factors did not significantly impact mid-fusiform activity (β = −0.33, p = 0.34). Further, to eliminate the confound of transition probabilities inherent to sentence construction, we analyzed activity for a word list condition (Figure S2). We found no significant interaction between word frequency and whether words were presented in a syntactically correct sentence or in an unstructured word list (β = −0.1, p = 0.87), disambiguating word frequency from predictability. We also assessed effects of other closely related parameters thought to be crucial to perceptual identification of words; bigram frequency and orthographic neighborhood for the pseudoword stimuli in mid-fusiform cortex. There were no significant effects of bigram frequency (β = 5.04, p = 0.08), mean positional bigram frequency (β = −0.14, p = 0.94) or orthographic neighborhood (β = 5.19, p = 0.30).

We also evaluated the effects of word frequency on the latency of lexical determination in mid-fusiform. We plotted the time course of activation in the mid-fusiform ROI to high, mid and low frequency words and pseudowords, matched for word length. This showed a clear association between word frequency and the latency of activity separation between words and pseudowords - high frequency words (180 ms) were distinguishable from pseudowords earliest, followed by mid-frequency words (270 ms) and finally low frequency words (400 ms) (Figure 4e).

### Temporal Dynamics of Lexical Processing

To replicate this analysis at the level of individual electrodes rather than a surface-based population analysis, we performed a multiple linear regression using broadband gamma activity at individual electrodes (Figure 4f). This, like the MEMA (Figure 4a), also revealed distinct separations in activity between mid-fusiform cortex and lateral occipitotemporal cortex. An LME model over time (Figure 4g) showed an effect of word length first at the occipital pole (75 ms) and then more anteriorly. Conversely, frequency sensitivity appeared earliest in lateral occipitotemporal cortex and mid-fusiform cortex (150 ms) and spread posteriorly.

Across these two tasks, we see two temporal stages of lexical selectivity; initial selectivity in mid-fusiform cortex followed by an anterior-to-posterior spread of selectivity. In our final analysis, we used an unsupervised clustering algorithm, non-negative matrix factorization (NNMF), on the time courses of the lexical and sub-lexical selectivity to quantify their spatiotemporal overlap and separability. An NNMF of the time course of d-prime selectivity of words vs all non-words in the sub-lexical task (Figure 5a) revealed a now familiar pattern - an early component in mid-fusiform and a late component in posterolateral vOTC and occipital sites (Figure 5d). These components were preserved when reanalyzed for each of the non-word conditions (Figure S3a,b). With increasing sub-lexical complexity, the early component diminished, and the late component remained highly consistent, representing latency differences in the ability of mid-fusiform to distinguish these conditions from words (Figure 3d; Figure S3c).

**Figure 5:**
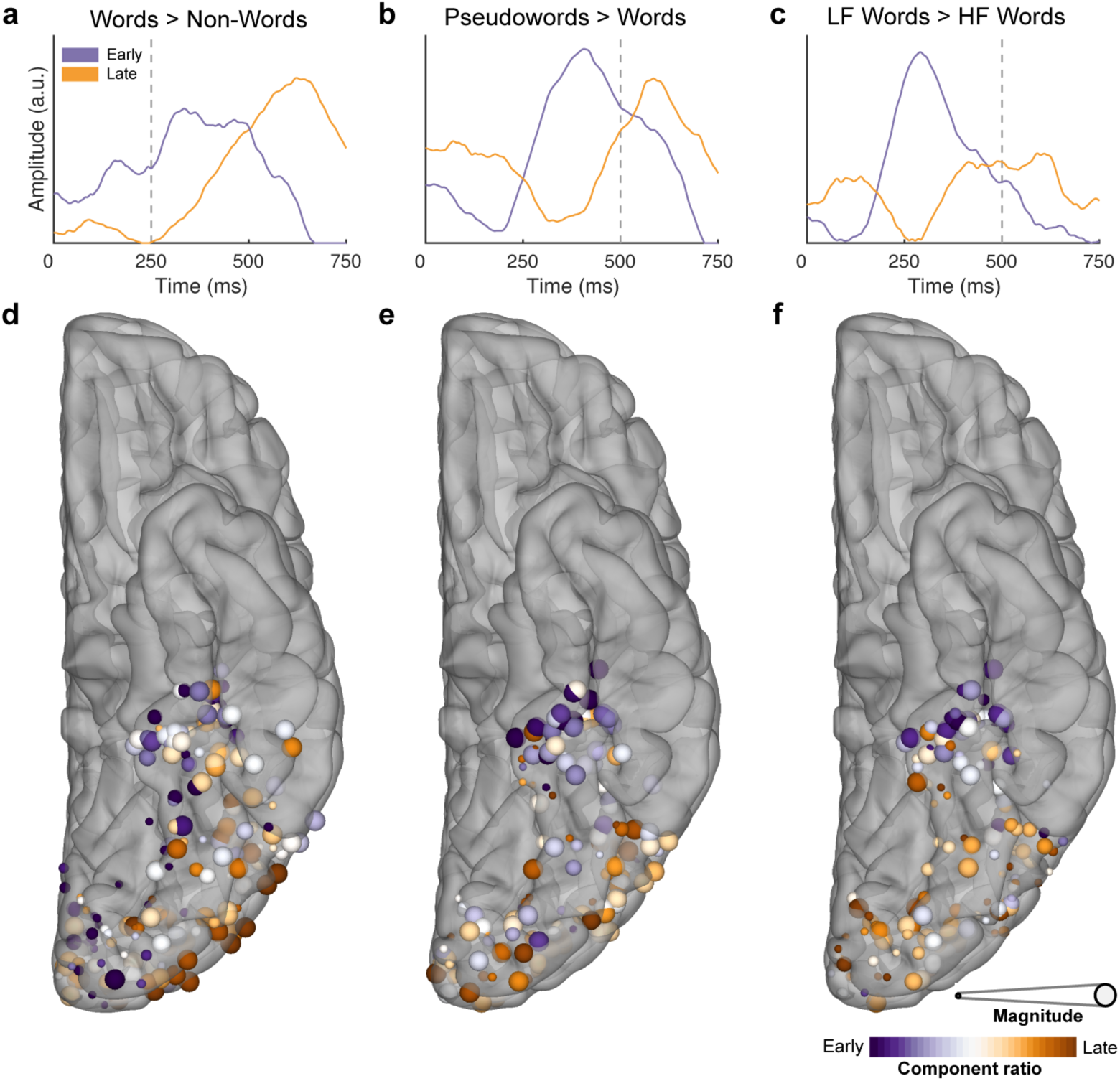
Antero-posterior differences in the timing of frequency and lexicality effects. Temporal (a,b,c) and spatial (d,e,f) representations of the two archetypal components generated from the NNMF for the contrasts of words vs non-words in the sub-lexical task (a,d), words vs pseudowords (b,e) and high (HF) vs low (LF) frequency words (c,f) during sentence reading. Vertical dashed lines denote word offset time. Spatial representations (d,e,f) are colored based on the weighting of their membership to either component. Size is based on the magnitude of the contrast between experimental conditions.

An NNMF analysis of the time courses of z-scores of lexical selectivity, distinguishing pseudowords from real words and high from low frequency words (Figure 5b,c), revealed an almost identical pattern - an early component in mid-fusiform cortex and a late component over posterolateral vOTC and occipital sites (Figure 5e,f). The late component was remarkably similar in time course to that seen in the sub-lexical task, however, the early component was variable across these two conditions, reflecting the differences in latency in the distinction of different frequency words discussed earlier.

## Discussion

Our work reveals two spatiotemporally distinct constituents of the vOTC that perform distinct roles in reading: the mid-fusiform cortex and lateral occipitotemporal gyrus. The amplitude and latency of the activity in these regions is highly sensitive to the statistics of natural language. This work shows the central role of the mid-fusiform cortex in both word vs. non-word discrimination and lexical identification.

That orthographic-to-lexical transformation occurs in mid-fusiform cortex is concordant with emerging evidence from lesion-symptom mapping (Pflugshaupt et al., 2009; Rodríguez-López et al., 2018; Tsapkini and Rapp, 2010) and intracranial stimulation studies (Hirshorn et al., 2016; Mani et al., 2008) of orthographic coding. We find unambiguously that the lateral occipitotemporal cortex, while active earlier, displays sensitivity to orthographic and lexical processing later than mid-fusiform cortex. The early and late selectivity in these regions during both sub-lexical and lexical processing is a potential correlate of bottom-up and top-down processes. This existence of two separable ventral cortical regions with distinct lexical and sub-lexical sensitivities is concordant with the view of a non-unitary VWFA (Bouhali et al., 2019; Lerma-Usabiaga et al., 2018; White et al., 2019).

An orthographic lexicon (Coltheart, 2004; Coltheart et al., 2001), or the long-term memory representations of which letter strings correspond to familiar words would be tuned to word frequency and lexicality. These two features were found to be coded earliest in this region and drive its activity – our model, incorporating just word frequency and length explained 73% of the variance of mid-fusiform activation. This central role of the mid-fusiform has also recently been suggested by selective hemodynamic changes following training to incorporate new words into the lexicon (Glezer et al., 2015; Taylor et al., 2019).

The latency distinctions in the mid-fusiform for words of varying lexical and sub-lexical frequency are consistent with a heuristic for searching the lexicon on the basis of word frequency and stopping the search once a match is found (Gold and Shadlen, 2004; Norris, 2006). Under this hypothesis, higher frequency words are matched to a long-term memory representation faster than infrequent words, and pseudowords require the longest search times, since they do not match any long-term memory representations. These latency differences invoke a possible role for this region as the “*bottleneck*” that limits reading speed (Rayner, 1977; Rayner and Duffy, 1986; White et al., 2019).

This work validates functional imaging studies of sensitivity to word frequency (Graves et al., 2010; Kronbichler et al., 2004; Schuster et al., 2016; White et al., 2019) and of greater activation to attended pseudowords than known words in the vOTC (Kronbichler et al., 2004). However, in contrast to previous fMRI studies, we found no evidence for a gradual buildup of sub-lexical complexity along the anteroposterior axis. Instead, there was a sharp transition between posterior regions and mid-fusiform cortex, where we see latency differences in processing of increasingly word-like letter strings followed by an anterior-to-posterior spread of lexical selectivity. Based on the hierarchy of word-likeness tested with the stimuli, the largest distinction was between the IL condition and the FL condition. One possibility for this distinction between these two classes of stimuli was that the IL condition consisted purely of consonant strings, while 75% of strings in the FL condition contained at least one vowel – a minimal requirement for plausibility in orthographic processing. Precisely which properties of letter sequence dictate the discrete transition between plausible and implausible words is an area of ongoing investigation.

Given previous behavioral and imaging results, we initially predicted sensitivity to orthographic neighborhood or bigram frequency in this region, both of which have previously been shown to influence speed and accuracy of non-word identification (Carreiras et al., 1997; Grainger et al., 2012; Rice and Robinson, 1975). During passive viewing we observed latency differences in word/non-word discrimination in mid-fusiform based on n-gram frequencies. However, during sentence reading neither of these factors showed significant effects on pseudoword activation in mid-fusiform cortex. The influence of these factors on orthographic processing may depend on the demands of the specific task (Meade et al., 2019). Specifically, both factors have been shown to play a role in how quickly participants reject non-words in lexical decision, and may be more indicative of how participants perform that particular task rather than reflect the automatic word identification processes.

The existence of an anterior-to-posterior spread of lexical and sub-lexical information from mid-fusiform cortex to earlier visual processing regions implies recursive feedback and feedforward interactions between multiple stages of visual processing within the ventral stream. This notion has a storied past in cognitive models of reading, including the interactive activation model (McClelland and Rumelhart, 1981), its derivatives (Coltheart et al., 2001; Perry et al., 2007), and the interactive account (Price and Devlin, 2003, 2011). The direct measurement of this anterior-to-posterior spread from mid-fusiform implies its role in mediating input from frontal regions during word (Whaley et al., 2016; Woodhead et al., 2014) and object (Bar et al., 2006) recognition, as predicted by others (Lerma-Usabiaga et al., 2018).

While the involvement of mid-fusiform in aspects of both sub-lexical and lexical processing in reading is reasonably unambiguous, the specificity of this region to orthographic input needs more study, perhaps at scales smaller than afforded by the electrodes used here. We have previously shown that the left mid-fusiform cortex is a critical lexical hub for both visually and auditory cued naming (Conner et al., 2014; Forseth et al., 2018), and these data imply that it is in fact a multi-modal lexical hub whose role includes encoding orthographic information. However, stimulation (Hirshorn et al., 2016; Mani et al., 2008) or lesioning (Pflugshaupt et al., 2009; Rodríguez-López et al., 2018; Tsapkini and Rapp, 2010) of the mid-fusiform can lead to selective disruption of orthographic naming, potentially suggesting separable orthographic specific regions in the mid-fusiform. This could also be interpreted as there being a lack of redundant processing pathways for written language as compared to other domains, resulting in orthographic processing being more susceptible to disruption.

In summary, we have demonstrated a central role of the mid-fusiform cortex in the early processing of the statistics of lexical and sub-lexical information in visual word reading. We have characterized the activity of mid-fusiform cortex as being sensitive, in both amplitude and latency, to the frequencies of occurrence of words in natural language. Further, we have shown the existence of an anterior-to-posterior spread of lexical information from mid-fusiform to earlier visual regions including classical VWFA.

## Materials and Methods

### Participants

A total of 35 participants (17 male, 19-60 years, 5 left-handed, IQ 94 ± 13, age of epilepsy onset 19 ± 10 years) took part in the experiments after written informed consent was obtained. All experimental procedures were reviewed and approved by the Committee for the Protection of Human Subjects (CPHS) of the University of Texas Health Science Center at Houston as Protocol Number HSC-MS-06-0385. Inclusion criteria for this study were that the participants were English native speakers, left hemisphere dominant for language and did not have a significant additional neurological history (e.g. previous resections, MR imaging abnormalities such as malformations or hypoplasia). Three additional participants were tested but were later excluded from the main analysis as they were determined to be right hemisphere language dominant. Hemispheric dominance for language was determined either by fMRI activation (n = 1) or intra-carotid sodium amobarbital injection (n = 2).

### Electrode Implantation and Data Recording

Data were acquired from either subdural grid electrodes (SDEs; 7 patients) or stereotactically placed depth electrodes (sEEGs; 28 patients) implanted for clinical purposes of seizure localization of pharmaco-resistant epilepsy.

SDEs were subdural platinum-iridium electrodes embedded in a silicone elastomer sheet (PMT Corporation; top-hat design; 3mm diameter cortical contact) and were surgically implanted via a craniotomy following previously described methods (Conner et al., 2011; Pieters et al., 2013; Tandon, 2012).

sEEG probes (PMT corporation, Chanhassen, Minnesota) were 0.8 mm in diameter, had 8-16 contacts and were implanted using a Robotic Surgical Assistant (ROSA; Medtech, Montpellier, France) (Tandon et al., 2019). Each contact was a platinum-iridium cylinder, 2.0 mm in length with a center-to-center separation of 3.5-4.43 mm. Each patient had multiple (12-20) probes implanted.

Following implantation, electrodes were localized by co-registration of pre-operative anatomical 3T MRI and post-operative CT scans using a cost function in AFNI (Cox, 1996). Electrode positions were projected onto a cortical surface model generated in FreeSurfer (Dale et al., 1999), and displayed on the cortical surface model for visualization (Pieters et al., 2013).

Intracranial data were collected using the NeuroPort recording system (Blackrock Microsystems, Salt Lake City, Utah), digitized at 2 kHz. They were imported into MATLAB initially referenced to the white matter channel used as a reference by the clinical acquisition system, visually inspected for line noise, artifacts and epileptic activity. Electrodes with excessive line noise or localized to sites of seizure onset were excluded. Each electrode was re-referenced offline to the common average of the remaining channels. Trials contaminated by inter-ictal epileptic spikes were discarded.

### Stimuli and Experimental Design

27 participants undertook a sub-lexical processing task and 28 participants undertook a rapid serial visual presentation (RSVP) sentence reading task, reading real sentences, Jabberwocky sentences and word lists.

All stimuli were displayed on a 15.4” 2880×1800 LCD screen positioned at eye-level at a distance of 80 cm and presented using Psychtoolbox (Kleiner et al., 2007) in MATLAB.

#### Sub-Lexical Processing

Participants were presented with 80 runs, each six stimuli in length and containing one six-character stimulus from each of six categories in a pseudorandom order. Stimulus categories, in increasing order of sub-lexical structure, were (1) false font strings, (2) infrequent letters, (3) frequent letters, (4) frequent bigrams, (5) frequent quadrigrams and (6) words (Figure 1a). n-gram frequencies were calculated from the English Lexicon Project (Balota et al., 2007). False fonts used a custom-designed pseudo-font with fixed character spacing. Each letter was replaced by an unfamiliar shape with an almost equal number of strokes and angles and similar overall visual appearance. The stimuli were based on a previous study (Vinckier et al., 2007), converted for American English readers.

A 1500 ms fixation cross was presented between each run. During each run, each stimulus was presented for 250 ms followed by a blank screen for 500 ms. Words were presented in all capital letters in Arial font with a height of 150 pixels. To maintain attention participants were tasked to press a button on seeing a target string of ###### presented. The target stimulus was inserted randomly into 20 runs as an additional stimulus and was excluded from analysis. Detection rate of the target stimuli was 91 ± 10 %.

#### Sentence Reading

Participants were presented with eight-word sentences using an RSVP format (Figure 1c). A 1000 ms fixation cross was presented followed by each word presented one at a time, each for 500 ms. Words were presented in all capital letters in Arial font with a height of 150 pixels. To maintain the participants’ attention, after each sentence they were presented with a two alternative forced choice, deciding which of two presented words was present in the preceding sentence, responding via a key press. Only trials with a correct response were used for analysis. Overall performance in this task was 92 ± 4 % with a response time of 2142 ± 782 ms.

Stimuli were presented in blocks containing 40 real sentences, 20 Jabberwocky sentences and 20 word lists in a pseudorandom order. Each participant completed between 2-4 blocks.

Word choice was based on stimuli used for a previous study (Fedorenko et al., 2016). Jabberwocky words were selected as pronounceable pseudowords, designed to fill the syntactic role of nouns, verbs and adjectives by inclusion of relevant functional morphemes.

### Signal Analysis

A total of 5666 electrode contacts were implanted, 891 of these were excluded from analysis due to proximity to the seizure onset zone, excessive interictal spikes or line noise.

Electrode level analysis was limited to a region of interest (ROI) based on a brain parcellation from the Human Connectome Project (Glasser et al., 2016). The ROI encompassed all areas deemed to be visually responsive in the Glasser atlas, including the entire occipital lobe and most of the ventral temporal surface, excluding parahippocampal and entorhinal regions (Figure 2b).

Analyses were performed by first bandpass filtering raw data of each electrode into broadband gamma activity (BGA; 70-150Hz) following removal of line noise (zero-phase 2nd order Butterworth bandstop filters). A frequency domain bandpass Hilbert transform (paired sigmoid flanks with half-width 1.5 Hz) was applied and the analytic amplitude was smoothed (Savitzky - Golay FIR, 3rd order, frame length of 151 ms; Matlab 2017a, Mathworks, Natick, MA). BGA is presented here as percentage change from baseline level, defined as the period −500 to −100 ms before each run in the sub-lexical task or before word 1 of each sentence.

Electrodes were tested to determine whether they were word responsive within the window 100-400ms post stimulus onset, a time window previously used for determining selectivity of vOTC (Kadipasaoglu et al., 2016). This was done by measuring the response of real words in the sub-lexical processing task, or all words in word position 1 in sentence reading. Responsiveness threshold was set at 20% amplitude increase above baseline with p<0.01. 601 and 459 electrodes respectively were located in left, language-dominant vOTC for the sub-lexical and lexical tasks of which 211 and 196 (in 20 patients each) were word responsive (Figure 2b).

When presented as grouped electrode response plots, within-subject averages were taken of all electrodes within each ROI then presented as the across subject average, with colored patches representing ±1 standard error.

### Linguistic Analysis

When separating content and function words, function words were defined as either articles, pronouns, auxiliary verbs, conjunctions, prepositions or particles. We quantified word frequency as the base-10 log of the SUBTLEXus frequency (Brysbaert and New, 2009). This resulted in a frequency of 1 meaning 10 instances per million words and 4 meaning 10,000 instances per million words. Bigram frequency was calculated as the mean frequency of each adjacent two letter pair, as calculated from the English Lexicon Project (Balota et al., 2007). Orthographic neighborhood was quantified as the orthographic Levenshtein distance (OLD20); the mean number of single character edits required to convert the word into its 20 nearest neighbors (Yarkoni et al., 2008).

### Statistical Modelling

#### Word Selectivity

Determination of the onset time of word selectivity within individual electrodes was determined as the first time point where the d-prime of the words against all non-word stimuli became significant at p<0.01 for at least 50 ms. The significance threshold was determined by bootstrapping the results with randomly assigned category labels, using 1000 repetitions.

#### Non-negative matrix factorization (NNMF)

NNMF is an unsupervised clustering algorithm (Berry et al., 2007). This method expresses a non-negative matrix A as the product of “class weight” matrix W and “class archetype” matrix H, minimizing ||A – WH||^2^.

The factorization rank k = 2 was chosen for all analyses in this work. Repeat analyses with higher ranks did not identify additional response types. Inputs to the factorization were d-prime values (Figure 5a) or z-scores (Figure 5b,c, Figure S3a) that were half-wave rectified. These were calculated for the m electrodes at n time points for the temporal analyses. Factorization generated a pair of class weights for each electrode and a pair of class archetypes – the basis function for each class. Component ratio was defined as the magnitude normalized ratio between the class weights at each electrode. Magnitude was defined as the sum of class weights at each electrode.

#### Surface-based mixed-effects multilevel analysis (SB-MEMA)

SB-MEMA was used to provide statistically robust (Argall et al., 2006; Fischl et al., 1999; Saad and Reynolds, 2012) and topologically precise (Conner et al., 2014; Esposito et al., 2013; Forseth et al., 2018; Miller et al., 2007) effect estimates of band-limited power change from the baseline period. This method, developed and described previously by our group (Kadipasaoglu et al., 2014, 2015), accounts for sparse sampling, outlier inferences, as well as intra- and inter-subject variability to produce population maps of cortical activity. Significance levels were computed at a corrected alpha-level of 0.01 using family-wise error rate corrections for multiple comparisons. The minimum criterion for the family-wise error rate was determined by white-noise clustering analysis (Monte Carlo simulations, 1000 iterations) of data with the same dimension and smoothness as that analyzed (Kadipasaoglu et al., 2014). All maps were smoothed with a geodesic Gaussian smoothing filter (3 mm full-width at half-maximum) for visual presentation.

Amplitude normalized maps were created by normalizing to the beta values of an activation mask. The activation mask comprised of significant activation clusters satisfying the following conditions; corrected p<0.01, beta>10% and coverage>2 patients.

To produce the activation movies, SB-MEMA was run on short, overlapping time windows (150 ms width, 10 ms spacing) to generate the frames of movies portraying cortical activity.

### Linear Mixed Effects (LME) Modelling

For grouped electrode statistical tests, a linear mixed effects model was used. LME models are an extension on a multiple linear regression, incorporating fixed effects for fixed experimental variables and random effects for uncontrolled variables. The fixed effects in our model were word length and word frequency and our random effect was the patient. Word length refers to the number of letters in the presented word. This variable was mean-centered to avoid an intercept at an unattainable value, namely a zero-letter word. Word frequency was converted to an ordinal variable to facilitate combination across patients. The ordinal categories for frequency (*f*) were very high (*f*>3.5), high (2.5< *f* ≤3.5), mid (1.5< *f* ≤2.5), low (0.5< *f* ≤1.5) and very low (*f*≤0.5). The random effect of patient allowed a random intercept for each patient to account for differences in mean response size between patients.

These predictors were used to model the average BGA in the window 100-400ms after word onset. Word responses within each length/frequency combination were averaged within patient. Patients only contributed responses to length/frequency combinations for which they had at least five word-epochs to be averaged together.

For single electrode analysis of the frequency effect a multiple linear regression was used. Factors word length and word frequency were again used. Word length was again mean-centered. Word frequency was treated as a continuous variable. For this analysis all the word epochs from the sentence and word list conditions were used. Results were corrected for multiple comparisons using a Benjamini-Hochberg False Detection Rate (FDR) threshold of q<0.05.

Time courses of length and frequency representation were tested using the LME model with 25 ms, non-overlapping windows. Significance was accepted at an FDR corrected threshold of q<0.01.

## Acknowledgements

The authors would like to thank Yujing Wang for assistance coordinating patient data transfers and Eliana Klier for comments on previous versions of this manuscript. We express our gratitude to all the patients who participated in this study; the neurologists at the Texas Comprehensive Epilepsy Program who participated in the care of these patients; and all the nurses and technicians in the Epilepsy Monitoring Unit at Memorial Hermann Hospital who helped make this research possible. This work was supported by the National Institute of Neurological Disorders and Stroke grant NS098981.

## Author Contributions

Conceptualization: OW, NT, SD; Methodology: OW, CD, NT, SD; Data curation: OW, CD, PSR; Software: OW, KJF, CD; Formal Analysis: OW; Writing – Original Draft: OW; Writing – Review and Editing: OW, NT, SFB, YL, SD; Visualization: OW; Supervision: NT; Project Administration: NT; Funding Acquisition: NT, SD, NEC.

## Supplementary

Video 1: **Spatiotemporal map of lexical sensitivity in ventral visual cortex**. MEMA activation video showing the regions of significant activation to the real word (W; left) stimuli and infrequent letter (IL; middle) stimuli. The word normalized amplitude map (right) shows regions with preferential activation to words (red) or infrequent letters (blue).

Video 2: **Spatiotemporal map of sensitivity to sub-lexical structure in ventral visual cortex**. MEMA video showing word normalized activation amplitudes for each of the non-word conditions from the sub-lexical task, demonstrating regions with preferential activation to words (red) or non-words (blue). FF: False Font, IL: Infrequent Letters, FL: Frequent Letters, BG: Frequent Bigrams, QG: Frequent Quadrigrams.

**Figure S1:**
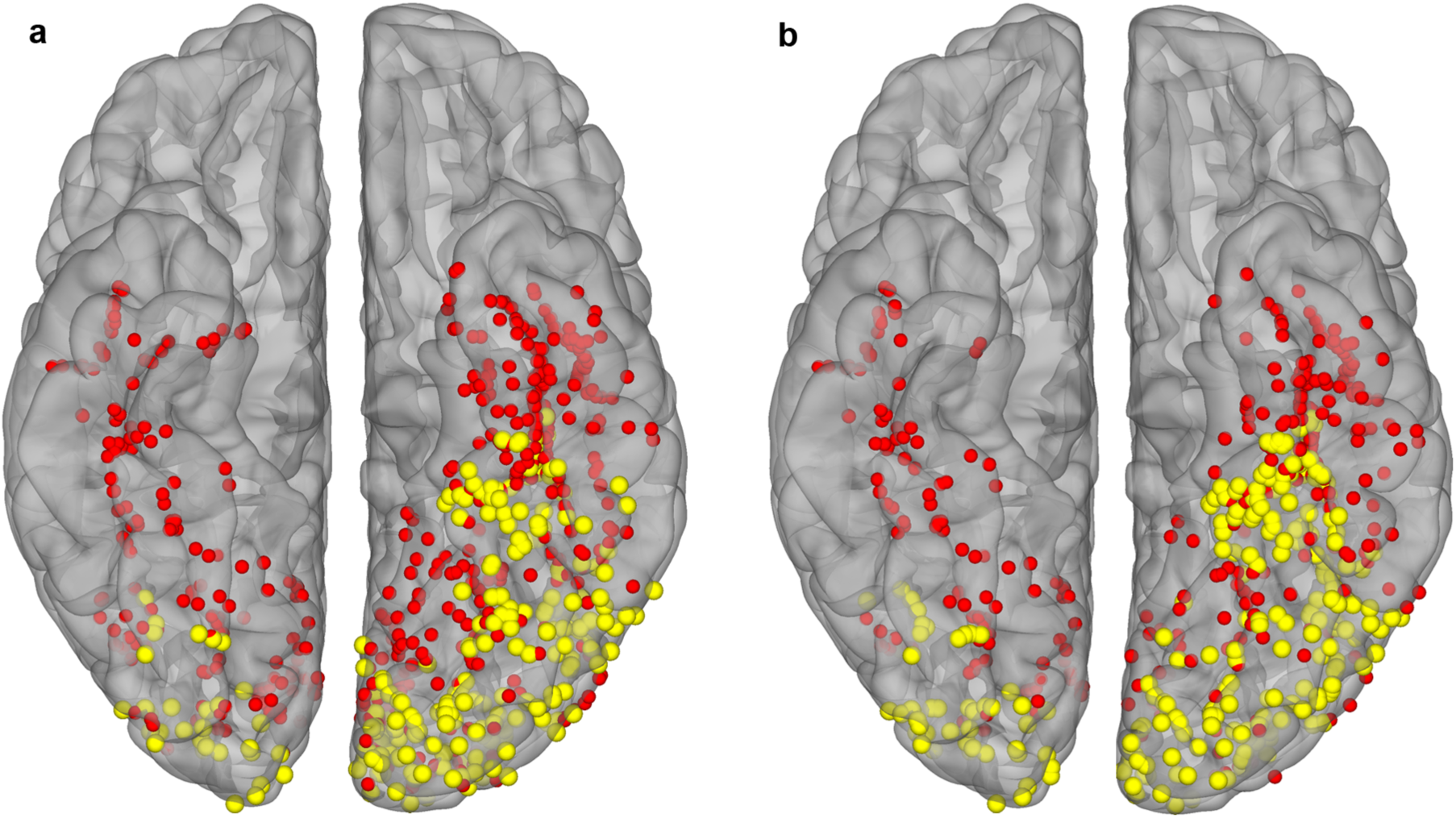
Lateralization of Word Responsive Electrodes in Ventral Cortex. Map of word responsive (yellow; activation >20% above baseline) and unresponsive (red) electrodes in the sub-lexical (a) and sentence (b) tasks. In the non-dominant right hemisphere (n = 14 patients), word responses were confined to occipital cortex.

**Figure S2:**
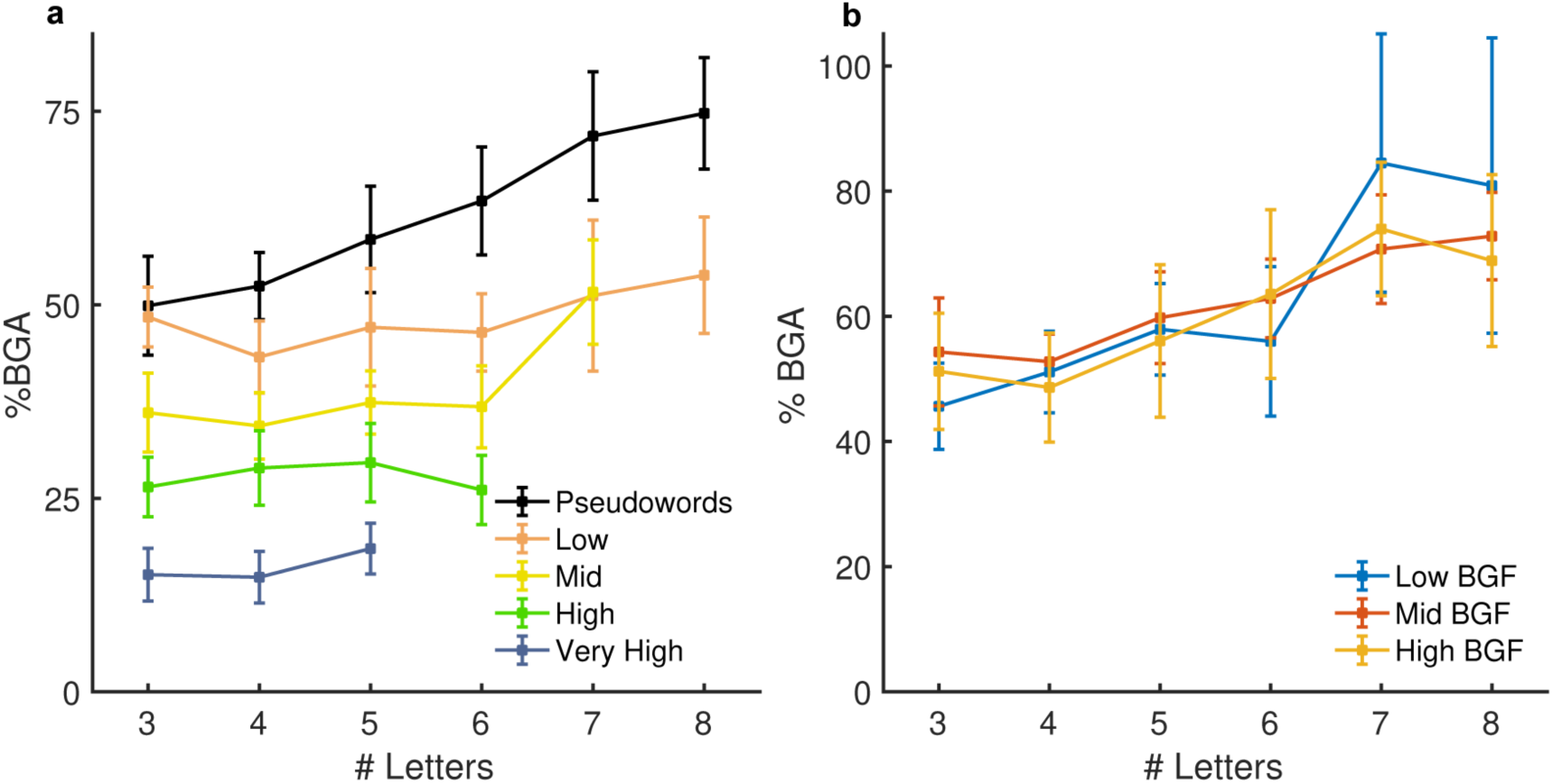
Lexical and Sub-Lexical Frequency Effects in Mid-Fusiform Cortex. (a) Mid-fusiform responses to real words from the word list condition separated by word frequency and length. (b) Pseudoword responses in mid-fusiform cortex from the Jabberwocky condition separated by bigram frequency (BGF) and word length.

**Figure S3:**
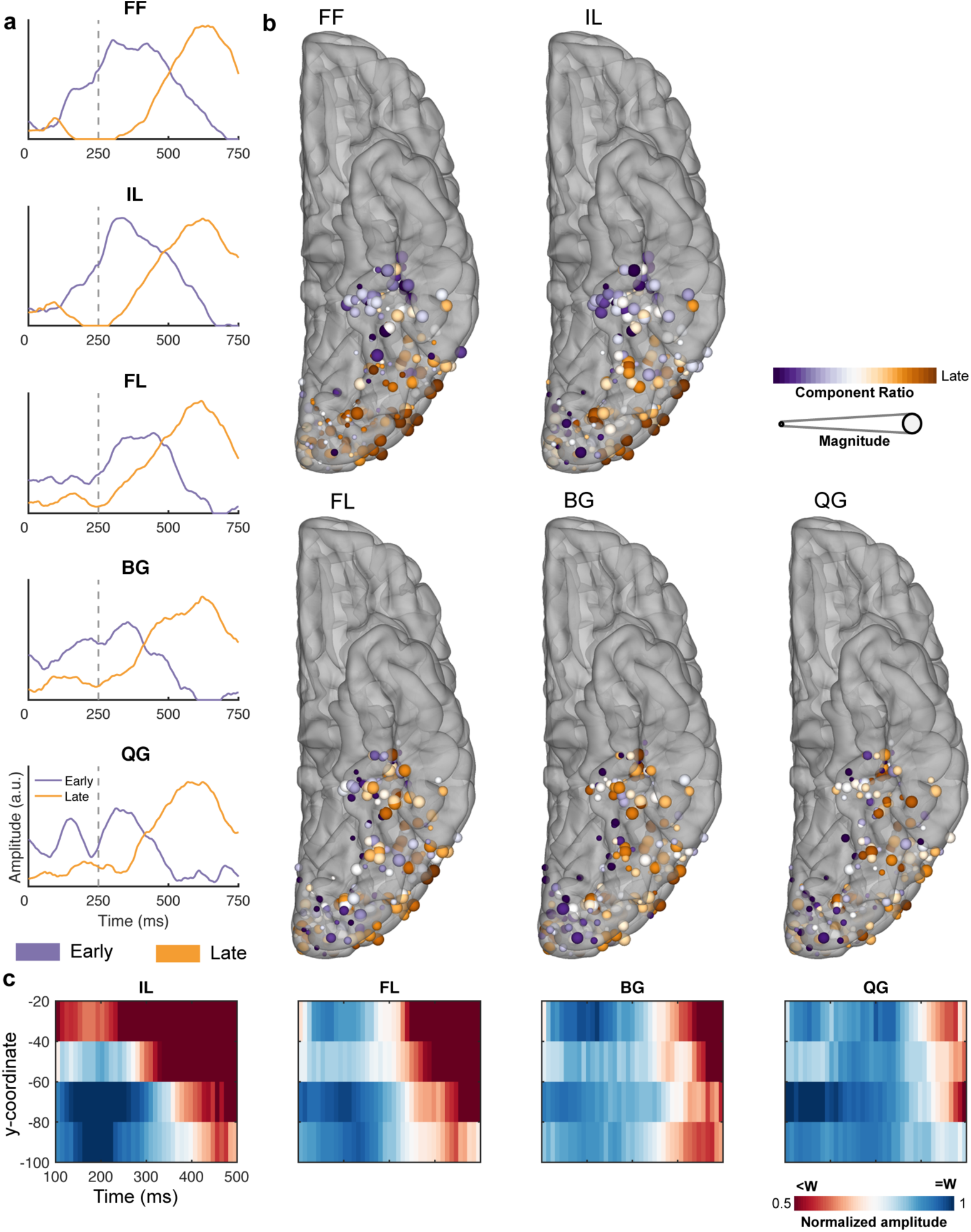
Timing of the selectivity to hierarchical orthographic stimuli in the sub-lexical task. (a) Temporal representations of the two archetypal components generated from each NNMF of the z-scores of words against each non-word condition. (b) Spatial map of the NNMF decompositions of the z-score word selectivity. (c) Spatiotemporal representation of word vs non-word selectivities (non-word normalized to word activity) for each of the letter-form conditions. Electrode selectivity profiles were grouped every 20 mm along the antero-posterior axis in Talairach space. Each condition shows an anterior-to-posterior spread of word selectivity (red). FF: False Font, IL: Infrequent Letters, FL: Frequent Letters, BG: Frequent Bigrams, QG: Frequent Quadrigrams.

